# The role of dectin-1 signaling in altering tumor immune microenvironment in the context of aging

**DOI:** 10.1101/2021.02.15.431361

**Authors:** Natarajan Bhaskaran, Sangeetha Jayaraman, Cheriese Quigley, Prerna Mamileti, Mahmoud Ghannoum, Aaron Weinberg, Jason Thuener, Quintin Pan, Pushpa Pandiyan

## Abstract

An increased accumulation of immune-dysfunction-associated CD4^+^Foxp3^+^ regulatory T cells (T_regs_) is observed in aging oral mucosa during infection. Here we studied the function of T_regs_ during oral cancer development in aging mucosa. First, we found heightened proportions of T_regs_ and myeloid-derived suppressor cells (MDSC) accumulating in mouse and human oral squamous cell carcinoma (OSCC) tissues. Using the mouse 4-Nitroquinoline 1-oxide(4-NQO) oral carcinogenesis model, we found that tongues of aged mice displayed increased propensity for epithelial cell dysplasia, hyperplasia, and accelerated OSCC development, which coincided with significantly increased abundance of IL-1β, T_regs_, and MDSC in tongues. Partial depletion of T_regs_ reduced tumor burden. Moreover, fungal abundance and dectin-1 signaling were elevated in aged mice suggesting a potential role for dectin-1 in modulating immune environment and tumor development. Confirming this tenet, dectin-1 deficient mice showed diminished IL-1β, reduced infiltration of T_regs_ and MDSC in the tongues, as well as slower progression and reduced severity of tumor burden. Taken together, these data identify an important role of dectin-1 signaling in establishing the intra-tumoral immunosuppressive milieu and promoting OSCC tumorigenesis in the context of aging.

## Introduction

Head and neck cancers (HNC) including OSCC are the sixth most common cancers, accounting for an estimated 657,000 new cases and more than 330,000 deaths globally each year. The aging population (age >65) has grown to 727 million in 2020 and will double to nearly 1.5 billion, roughly ~16% of the world population by 2050. A majority of all new cancers are diagnosed among elderly adults [1]. Therefore, the convergence of the aging population and a higher cancer incidence in this population will result in a significant rise in cancer. Aging triggers the immune system to be in a chronic hyper-inflammatory state known as inflammaging. An imbalance between pro- and anti-inflammatory mechanisms due to the combination of inflammaging and impaired immune-surveillance may contribute to increased susceptibility of elderly individuals to cancer [2–4]. However, precise cellular alterations in the tumor immune microenvironment (TIME) that contribute to tumor growth and progression in the elderly are not completely understood. HNC are among the most highly immune-cell infiltrated cancer types, highlighting the significance of immune cells in tumor development and progression blockade. Immune-suppressive cells including T_regs_, MDSC, and tumor-associated macrophages (TAM) that are important in immune tolerance, accumulate in tumors, and are thought to contribute to poor immunologic responses against tumors resulting in cancer immune evasion [5–14]. Depending on the tumor type, T_regs_ may constitute between 30 - 45% of CD4^+^ T cells in solid tumors of nonlymphoid origin, with OSCC cancers having the highest degrees of T_reg_ infiltration[15; 16]. T_regs_ and MDSC have been shown to contribute to tumor evasion and even hinder the success of α-PD1 cancer immunotherapy[17–20]. Thus there is considerable interest in understanding the involvement of these cells in many cancers[21–25]. However, molecular mechanisms that instruct intra-tumoral T_reg_ recruitment, proliferation, stability, or functions in the context of their interactions with anti-tumor CD8 cells and other immune-suppressive cells remain elusive. Understanding these mechanisms will lead to new combinatorial strategies in the face of PD-1/PD-L1 blockade resistance, leading to improved patient outcomes. Having previously linked some of the immunosenescent cytokines in T_reg_ accrual and aging previously [26], we now show that dectin-1 signaling is required for increased IL-1β, MDSC, and T_regs_ during carcinogenesis, and that this enhancement in aged mice contributes to the acceleration in tumor development in them.

## Results

### Aged mice show earlier and exacerbated dysplasia/hyperplasia, and OSCC development as well as heightened levels of immunosuppressive cells in the tongue

To obtain mechanistic insights into immunological changes driving OSCC and to define aging-dependent alterations in early immunological events during carcinogenesis, we used the 4-NQO oral carcinogenesis mouse model. This constitutes 16-weeks of 4-NQO administration in drinking water followed by another 6-weeks with regular drinking water (**Fig. 1A**). We compared young (3-4 months of age) and aged (20-24 months of age) C57BL/6 mice treated with 4-NQO. Control mice received the propylene glycol vehicle in drinking water. At indicated times, we examined the tongue pathology by immuno-histochemistry using established OSCC criteria (**Fig. 1B**). Thickened keratinized layer (Hyperkeratosis; HK), lesions showing enlarged nuclei with increased nuclear to cytoplasmic ratio, loss of typical epithelial cell organization (dysplasia), intrusion into the connective tissue, and presence of cell nests were used for scoring on a 0-5 scale. Aged mice demonstrated earlier and higher incidence of accelerated progression of hyperkeratosis, hyperplasia, and dysplasia during early time points (**12-22 weeks**) (**Fig. 1B**). Also, more mice in the aged group showed these changes at early time-points (**Fig. S1**). These data are consistent with a previous study that elegantly showed similar results in aged mice [27]. Since aging-related differences in OSCC were not significant after 23-24 weeks, we focused on early events of carcinogenesis in defining altered and accelerated dysplasia in aging mice. Therefore, we examined the rest of the parameters at this early time (12-16 weeks) window. We isolated the tongue/gingival tissues and the oral draining cervical lymph nodes (CLN) and determined the frequency of CD3^+^CD4^+^Foxp3^+^ T_regs_ by flow cytometry (n=6/ Veh. group; n=8/ 4-NQO group). 4-NQO carcinogenesis process also increased the frequency of Foxp3^+^ T_regs_ in oral tissues and draining lymph nodes, which increased with time(**Fig. 1C, S2**). Aged mice displayed significantly higher proportions of T_regs_ when compared to younger mice (**Fig. 1C**). Correlating with the increase in T_regs_, the percentage of CD3^+^CD8^+^ cells decreased, and the aged mice had a significantly higher T_reg_: CD8 ratio than young mice (**Fig. 1D, left**). CD8^+^ T cells in the tongue also upregulated PD-1 but showed a decreased expression of IFN-γ in 4-NQO treated mice in both groups (**Fig. 1D, right**). Also, tongues from 4-NQO treated mice showed increased accumulation of CD11b^+^Ly6G^+^MDSC cells that were at higher levels in aged mice than young mice (**Fig. 1E**). Consistent with their MDSC phenotype, CD11b^+^Ly6G^+^ cells were Arginase-1^high^ (Arg-1), when compared to CD11b^+^Ly6G^-^ cells (**Fig. 1F, top**). Moreover, the expression of Arg-1 was much more elevated in aged 4-NQO mice than young 4-NQO mice (**Fig. 1F, bottom**). Thus, earlier epithelial dysplasia coincided with an increased T_regs_: CD8 ratio and CD11b^+^Ly6G^+^ Arg-1^high^ MDSC in aged tongues. Taken together, these results show that aging in the context of the carcinogen 4NQO, causes premature development of oral carcinogenesis, which is associated with increased infiltration of immune-suppressive cells.

**Fig. 1.**
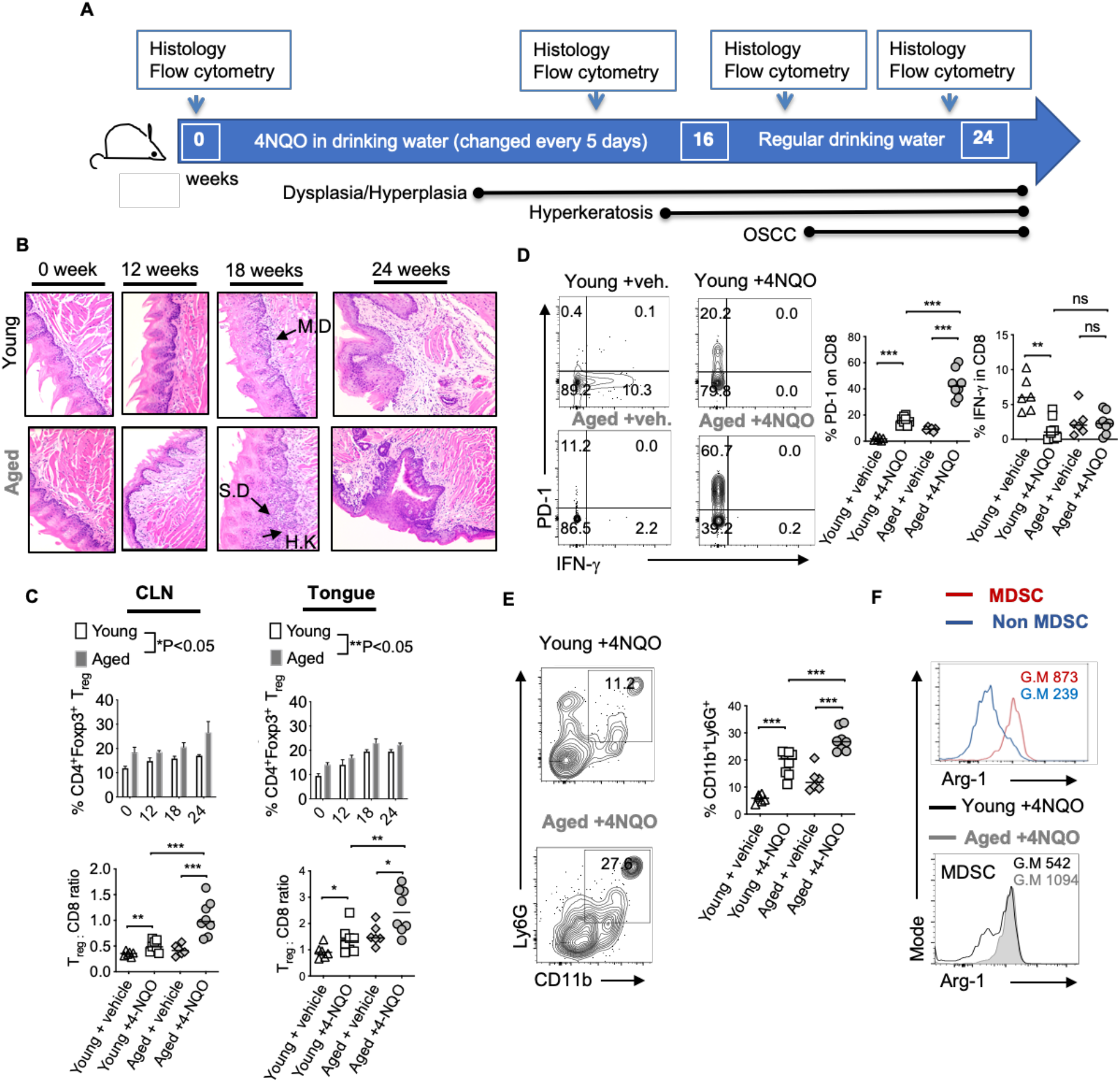
Aged mice display accelerated development of OSCC and heightened levels of immunosuppressive cells in the tongue. 4-NQO was (50 μg/ml) administered in drinking water. Time-points for readouts and various stages of OSCC development in this model are shown **(A)**. H&E immunohistochemistry images of the tongue at 200X magnification at indicated times [IF: Infiltration, MD: mild dysplasia, SD: severe dysplasia/hyperplasia, HK: Hyperkeratosis]. Samples were processed for flow cytometry **(B)**. Statistics showing %T_regs_ (**C**, top) and T_reg_:CD8 ratio (**C**, bottom) in the draining cervical lymph nodes (CLN) (left), or tongue (right) at 16 weeks (**D**). Contour plots (left) and statistics (right) showing % of PD-1 and IFN-γ expressing CD8^+^ cells (E). Representative contour plots (left) and statistics (right) showing % of CD11b and Ly6G expressing CD45+ cells. Arg-1 expression in MDSC (CD45^+^CD11b^+^Ly6G^+^) vs non MDSC (CD45^+^CD11b^+^Ly6G^-neg^ cells (F, top) and in MDSC in young and aged 4-NQO treated mice (**F**, bottom) (Geometric Mean = G.M)(Mann-Whitney test * P<0.05).

### Heightened levels of intra-tumoral T_regs_, MDSC, and IL-1β in OSCC patients

To validate 4-NQO findings, we examined surgically excised human OSCC tumors, control biopsies that included tissue derived from a site 2-3 cm from the tumor margin, as well as tumor lesion cytobrushings and contralateral control tissue cytobrushings from patients (**Fig. 2A, S3**). Tumor tissues were obtained from 16 patients that included eight former smokers, four current smokers, and four non-smokers. Similar to 4-NQO treated mice, human OSCC tumors showed very high proportions of CD25^+^Foxp3^+^ T_reg_ cells, (sometimes up to 42%; Mean 22%) (**Fig. 2B**). This was significantly higher than what is found in gingival mucosa of healthy individuals (*P<0.05)[26]. Among CD45^+^ cells, there were also higher proportions of CD14^neg^CD11B^+^CD33^+^ MDSC cells (**Fig. 2C**), which had higher ARG-1 expression, compared to non-MDSC cells(**Fig. 2D**). Because IL-1β signaling is critical for induction of immunomodulatory ROR-γt^+^FOXP3^+^ cells in oral mucosal tissues and is implicated in oral tumor progression[26; 28; 29], we examined the expression of IL-1β. IL-1β was significantly increased in the supernatants derived from human tumors and resected tumor supernatants (**Fig. 2E**), as well as in immune cells (**Fig. 2F**). However, the IL-1 receptor antagonist (IL-1RA) levels were significantly lower in tumor samples (**Fig. S4**) suggesting that IL-1 antagonism is associated with health while IL-1β was associated with tumor development. Notably, MDSC cells showed heightened levels of IL-1β than other cells, implying that IL-1β is likely linked to immune-suppressive milieu in tumors (**Fig. 2F**).

**Fig. 2.**
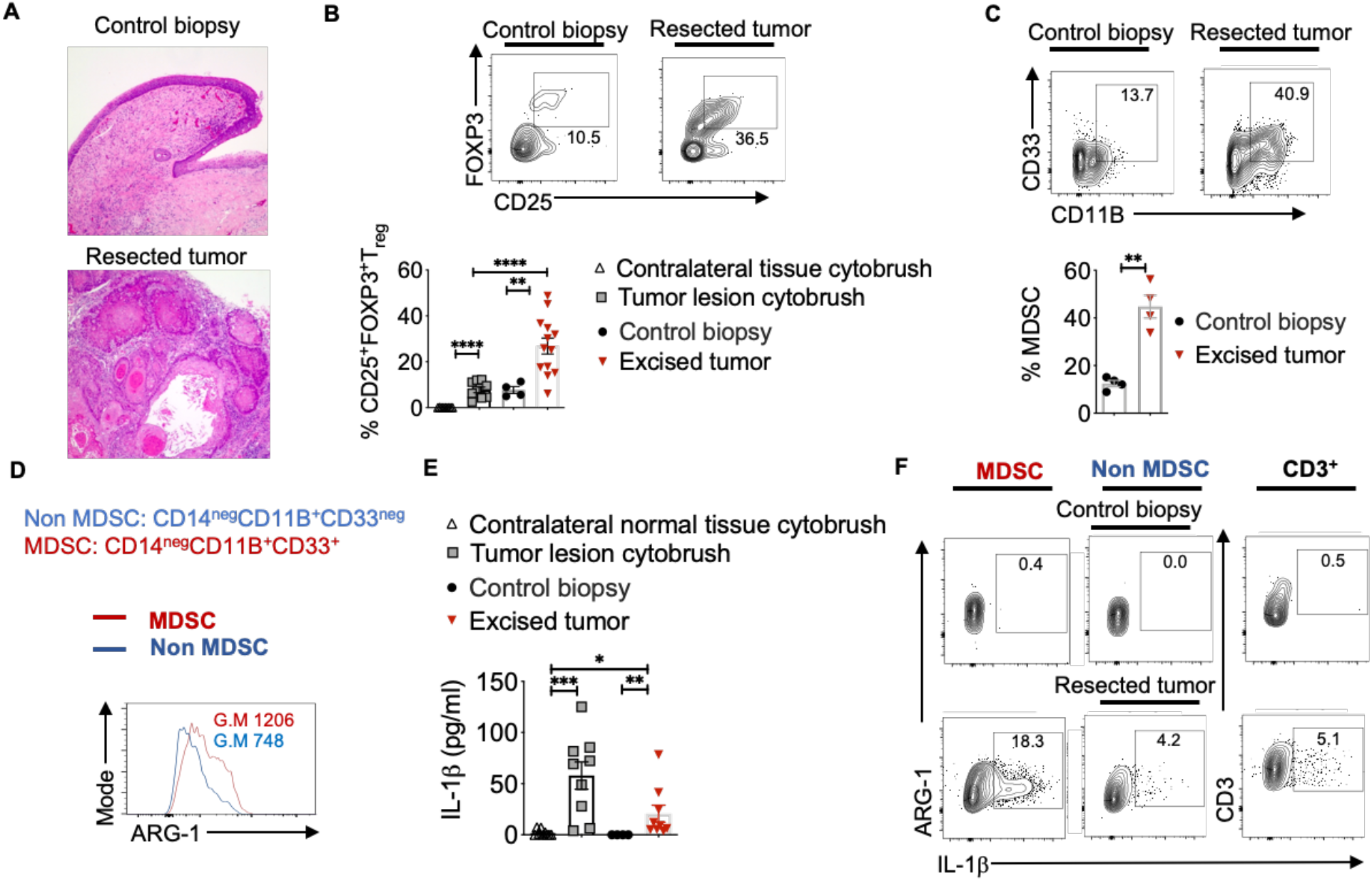
Heightened levels of intra-tumoral T_regs_, MDSC, and IL-1β in OSCC patients. Human oral tissue samples were obtained by excision (tumor resection), cytobrushing (mostly epithelial cells), or by biopsy (control tissue 2-3 cm from the tumor margin) under an approved IRB protocol. H&E immunohistochemistry images of the tongue at 200X magnification (**A**). Samples were digested by collagenase-1A and restimulated with PMA/Ionomycin for 4 hours. Contour plots gated on CD45^+^CD3^+^CD4^+^ lymphocytes (**B**, top) and statistical analysis (**B**, bottom) showing % of T_regs_ and MDSC (**C**). ARG-1 expression in MDSC vs non-MDSC cells (**D**). Supernatants were collected from these cultures and assessed for IL-1β using ELISA (**E**). Contour plots showing IL-1β expressing MDSC (left), non-MDSC (middle), and T cells (right) (**F**). Mean +/−SEM are shown. (Mann-Whitney test * P<0.05).

### Aged mice have a distinct T_reg_ phenotype and elevated levels of IL-1β compared to young mice

The degree of phenotypic and functional heterogeneity of tissue/intra-tumoral T_regs_ can be high, and evaluating them solely with FOXP3 and CD25 markers without examining anti-tumor CD8^+^ T cells can sometimes be misleading. Besides recruiting thymic T_regs_ into tumors via chemotaxis, T_regs_ can be induced *in situ* in tumors due to the abundance of mediators such as nitric oxide synthase, indoleamine 2, 3-dioxygenase 1, transforming growth factor-β1, and adenosine released from tumor cells, TAM, and MDSC [31]. Some of these FOXP3^+^ cells may also be critical for tissue repair or become dysfunctional depending on the inflammatory cytokine milieu[32]. Indeed we[26; 33–37] and others have shown distinct functions of T-bet^+^Foxp3^+^cells, ROR-γt^+^Foxp3^+^ cells, and PD-1^+^Foxp3^+^ cells, whose functions are significantly altered by cytokines including IL-6 and IL-1β in an mTOR dependent manner in oral mucosa[26]. Since CD25, PD-1, T-bet, ROR-γt, Suppression of Tumorigenicity 2 (ST-2) expression were all implicated in determining intra-tumoral T_reg_ functions positively and negatively in other tissues[7; 28; 38; 39], we examined the expression of the molecules, in aged vs. young T_regs_, 16 weeks after 4-NQO administration. While the expression of CD25 (ability to consume IL-2) [10; 40], Ki-67 (proliferation), and ST-2 (tissue T_reg_ and tumorigenicity marker) [7] did not change between young and aged T_regs_, CD103 (TGF-β1 dependent, tolerogenic, and tumor-specific T cell marker)[41], PD-1, ROR-γt, and T-bet were significantly up-regulated in aged T_regs_ (**Fig. 3A)**. Consistent with the higher expression of ROR-γt in T_regs_ and earlier dysplasia in aging mice, increased numbers of IL-1β expressing cells were found among infiltrating immune cells in aged mice as early as 5 and 12 weeks after 4-NQO administration (**Fig. 3B, C**). IL-6 levels were also elevated in aged tongue supernatants(**Fig. S5**). Moreover, at later times, IL-1β expression was found in epithelial tumor cells and immune cell infiltrates in both the groups, but was dramatically elevated in aged 4-NQO treated mice when compared to their young counterparts (**Fig. 3D**). These results showed that T_regs_ found in tongues during early tumor development in 4-NQO treated mice are phenotypically distinct, appearing to be functionally more immunosuppressive in aged mice, and may be linked to higher expression of IL-1β in the milieu.

**Fig. 3.**
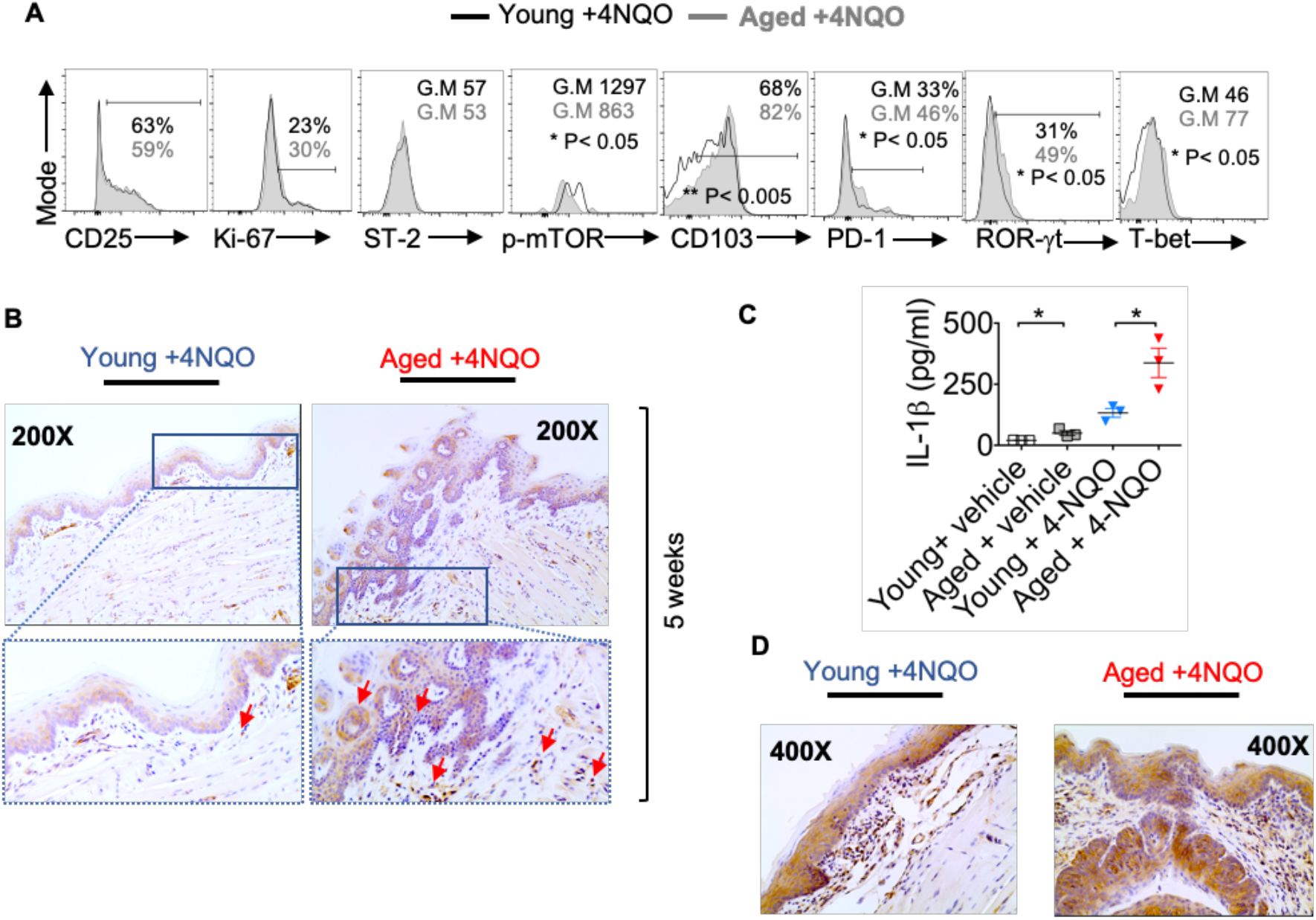
Aged mice have distinct T_reg_ phenotype and elevated levels of IL-1β compared to young mice. 16 weeks after 4-NQO administration, single-cell suspension from tongue was re-stimulated with PMA/Ionomycin for 4 hours. Flow cytometry histograms showing indicated protein expression in CD45^+^ CD3^+^ CD4^+^Foxp3^+^cells **(A)**. Tongue tissues were processed for IL-1β immuno-histochemical staining at 5 weeks(B). Supernatants collected at 12 weeks were processed for IL-1β ELISA. Mean +/− SEM in 3 mice/group (Mann-Whitney test * P< 0.05) (**C**). Tongue tissues were processed for IL-1β immuno-histochemical staining at 24 weeks(**D**).

### Partial T_reg_ depletion reduces tumor progression

Although tumorigenesis coincides with increases in Foxp3^+^ T_regs_, the function of these cells has not been validated in the 4-NQO model. Therefore, we employed Foxp3^DTR^-eGFP reporter (Foxp3 diphtheria toxin receptor; FDTR) mice that express the diphtheria toxin receptor on Foxp3^+^ cells and allow targeted T_reg_ depletion by intraperitoneal diphtheria toxin (DT) injection (20 μg/10 gm body weight; n=4/group)[36; 40]. We took two approaches; 1) Early and long-term T_reg_ depletion where DT was injected every 5 days, in the first 16 weeks of 4-NQO treatment 2) Late and shortterm T_reg_ depletion where DT was injected every 5 days, from the 16^th^ −21^st^ weeks of 4-NQO treatment n=6/group)(**Fig. 4A**). Early T_reg_ depletion caused systemic autoimmunity characterized by splenomegaly, lymphadenopathy, CD4 hyperactivation (CD44^high^, IFN-γ^high^), and mortality in 30% of mice which confounded the effects on early tumorigenesis (*not shown*). However, late and short-term T_reg_ depletion caused only partial T_reg_ reduction, but the mice had significantly reduced dysplasia and OSCC at 24 weeks (**Fig. 4B, C, S6A**). The frequencies of CD11b^+^Ly6G^+^ and IL-1β^+^ cells were unchanged with T_reg_ depletion, showing that these components are likely upstream to T_reg_ induction and infiltration during tumorigenesis (**Fig. S6B, C**). However, the PD-1 expression was diminished and IFN-γ expression was partially restored in CD8^+^ T cells(**Fig. 4D**). Collectively, these results show, for the first time in a 4-NQO OSCC model, that infiltrating T_regs_ control anti-tumorigenic functions of CD8^+^ T cells and contribute to OSCC tumorigenesis and progression.

**Fig. 4.**
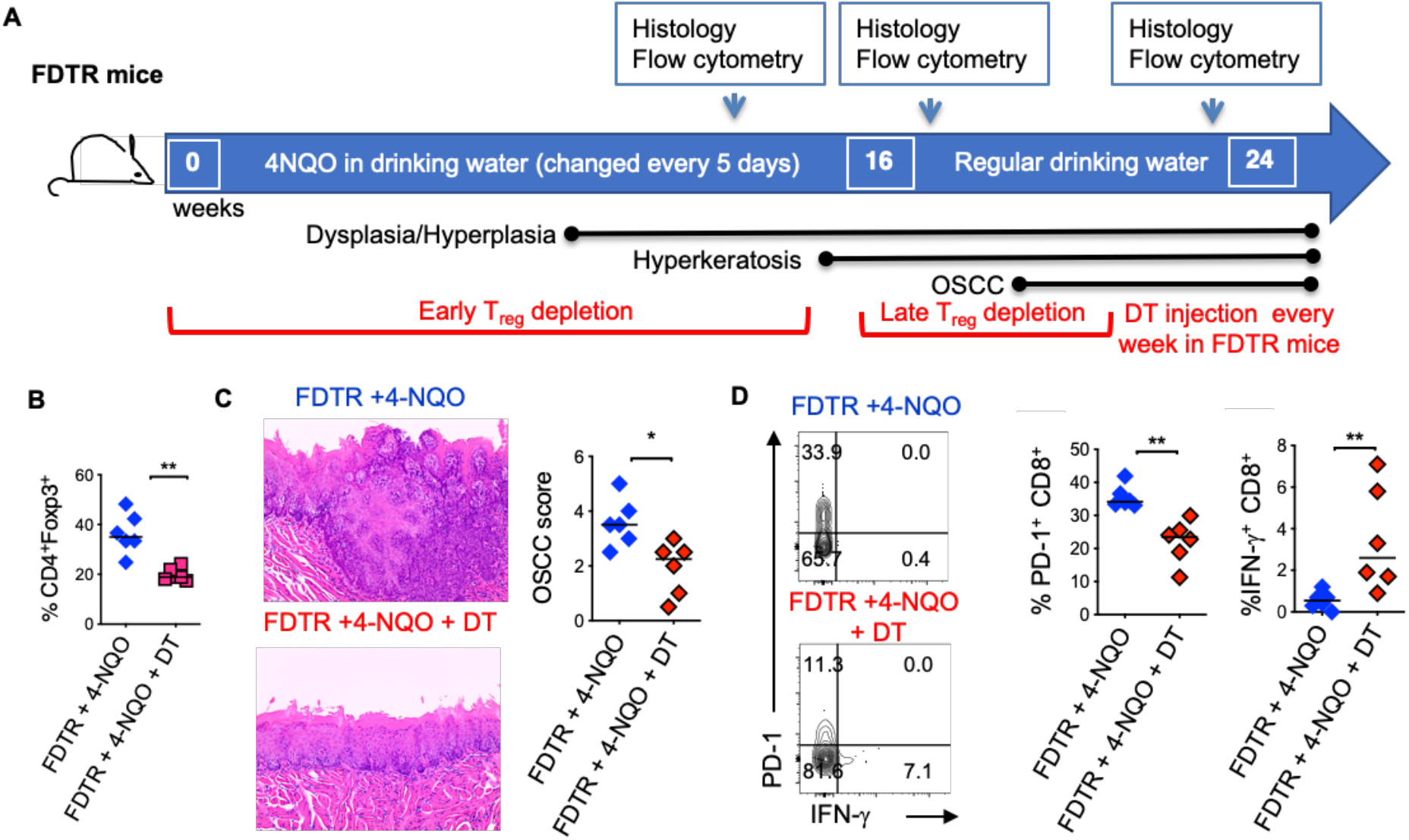
Partial T_reg_ depletion reduces tumor progression. T_regs_ were depleted in FDTR mice by injecting diphtheria toxin (DT) at late time points (every 5 days between 16^th^ −21^st^ weeks of 4-NQO treatment) **(A)**. Statistics showing CD4^+^Foxp3^+^cells based on flow cytometry analyses at 20 and 24 weeks **(B)**. Immunohistochemistry H&E staining (left) and OSCC score based on invasive hyperkeratosis, and nesting (0-5) (right) **(C)**. Contour plots (left) and statistics (right) showing PD-1 and IFN-γ expressing CD8^+^ T cells **(D)**. Mean +/− SEM in 6 mice/group (Mann-Whitney test * P< 0.05).

### *Candida* and zymosan exacerbate and accelerate dysplasia and hyperplasia

Previous evidence has shown that oral carriage of *Candida* is higher in elderly individuals and patients with OSCC[42–46]. We and others have previously shown that *Candida* infection can also induce IL-1β expression and T_reg_ infiltration in oral mucosa[26; 47]. Since aforementioned results showed IL-1β was positively linked with tumorigenesis, we hypothesized that *Candida* infection and consequent IL-1β induction might worsen OSCC outcome. Therefore, we examined the effect of *Candida* infection and zymosan during carcinogenesis in 4-NQO treated mice. We sublingually applied *Candida* (10^7^ blastospores) or D-zymosan (TLR-2 depleted zymosan; 1 mg/100ul) under anesthesia WT or TLR-2 knockout (TLR-2 KO) C57BL/6 mice every week between 4^th^ and 8^th^ weeks of 4-NQO administration. These treatments significantly accelerated and worsened dysplasia/hyperplasia (**Fig. 5A, B**), increased IL-1β^+^ cells and T_regs_ in the tongue at 12 weeks (**Fig. 5C, D, S7)**, and induced OSCC around 21 weeks (**Fig. 5E)**. Importantly, premature OSCC development was also observed in TLR-2 KO mice, revealing that zymosan and *Candida* mediated exacerbation of OSCC development was independent of TLR-2 signaling. In fact, in the absence of TLR-2, the numbers of IL-1β^+^ cells were increased implying the involvement of anti-fungal dectin-1 signaling in OSCC.

**Fig. 5.**
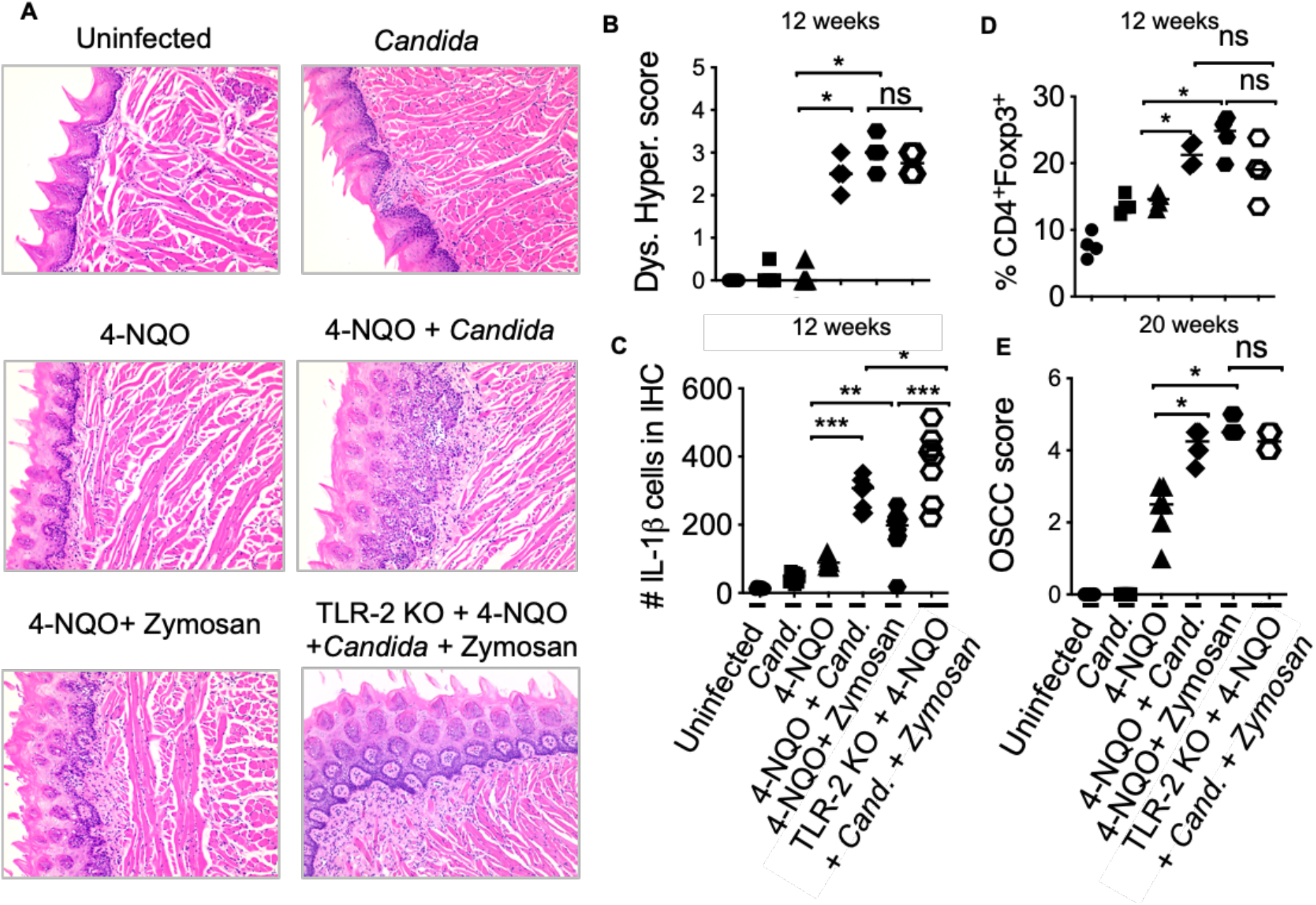
*Candida* and Zymosan exacerbate and accelerate dysplasia and hyperplasia. **A)** 4-NQO was administered in WT or TLR-2 KO C57BL/6 mice as in Fig. 3A (6 mice/group). Zymosan (1mg), *Candida* (10^7^ blastospores), or both were applied sublingually under anesthesia every week between the 4^th^ and 8^th^ weeks of 4-NQO administration. H&E staining was performed to assess dysplasia and hyperplasia (**A**). Immunohistochemistry and flow cytometry were performed. Statistics based on dysplasia and hyperplasia score (**B**), IL-1β immunohistochemistry (**C**), flow cytometry staining for T_reg_ assessment (**D**), and OSCC scores (**E**). OSCC scores were assigned based on hyperplasia, invasive hyperkeratosis, and nests after 20 weeks after 4-NQO administration. Statistics show Mean +/− SEM (Mann-Whitney test * P< 0.05).

### Aged mice show elevated Dectin-1 signaling in immune cells in the tongue and increased fungal abundance in saliva

Dectin-1 is expressed by myeloid phagocytes[48] and binds specifically to fungal β-1,3 glucans, endogenous galectins, and annexins on apoptotic cells[49]. Dectin1 signaling activation induces the phosphorylation of Syk and consequently the expression of IL-1β. We and others have shown that dectin-1 signaling can regulate T_reg_ alterations and macrophage metabolism[50–53]. Fungal dysbiosis and dectin-1 have also been implicated in inflammatory diseases and cancers[54–56]. Thus, previous reports and our above-mentioned results led us to hypothesize that dectin-1 might be involved in OSCC development and earlier predisposition of aged mice to OSCC. Confirming the hypothesis, 4-NQO induced tumorigenesis promoted a significant elevation of dectin-1 expression and Syk activation in immune infiltrates (CD45^+^ cells) (**Fig. 6A**). Dectin-1 expression and Syk activation were further increased in aged 4-NQO treated mice suggesting the involvement of dectin-1 signaling in accelerated tumorigenesis in aging mice(**Fig. 6B**). Although not-significant, there was also a trend towards increased fungal abundance in the saliva of 4-NQO treated mice, as measured by fungal abundance qPCR[57; 58]. Interestingly, aged 4-NQO treated mice that displayed accelerated dysplasia and OSCC (**Fig. 1A**), showed a significantly higher fungal abundance than control mice (**Fig. 6C**). Together, these results revealed that increased fungal abundance and resultant aberrant dectin-1 signaling might play a role in accelerated tumorigenesis in aged mice.

**Fig. 6.**
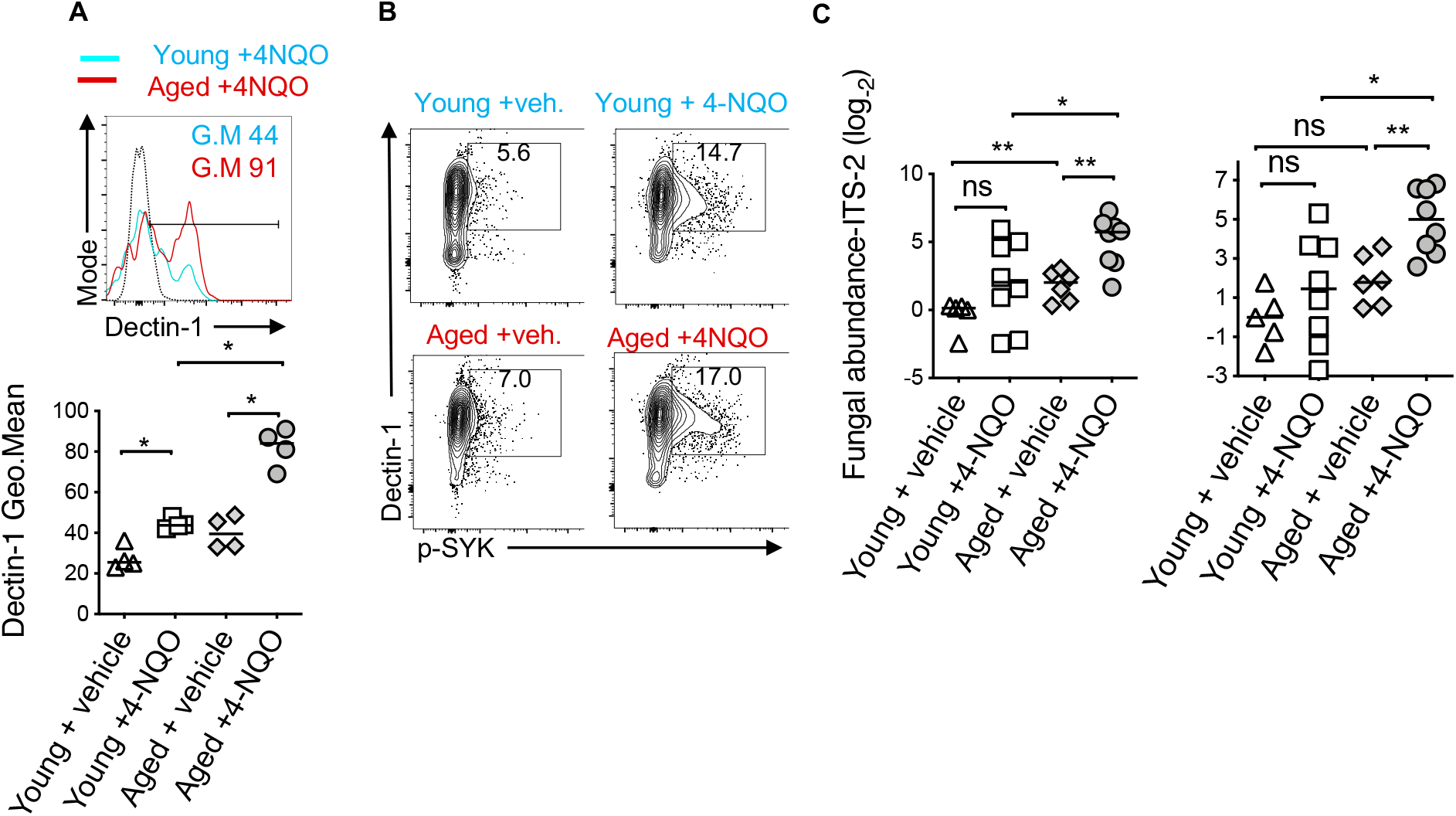
Aged mice show elevated dectin-1 signaling in immune cells in the tongue and increased fungal abundance in saliva. Tongue samples were processed for flow cytometry. Contour plots (top) and statistics (bottom) showing dectin-1 expressing CD45^+^ cells **(A)**. Contour plots showing % of Dectin-1^+^ and phosphorylated SYK (p-SYK) expressing CD45^+^ cells **(B)**. Saliva swabs collected at 12, 15, and 23 weeks were pooled. qPCR was performed to quantify fungal DNA using ITS-2 and fungi-quant PCR primers (**C**) [57; 58]. Relative abundance was standardized to input DNA using β-actin as reference. (Mann-Whitney test * P<0.05).

### Dectin-1 deficiency reduces immunosuppressive cells and OSCC tumor burden

Next, we investigated the role of dectin-1 in tumorigenesis using the mice deficient in dectin-1 signaling (Dectin KO mice). By monitoring body-weight we found that weight loss was much less pronounced in Dectin-1 KO than WT mice (**Fig. 7A**). Dectin-1 KO mice also had significantly reduced IL-1β^+^ cells (**Fig. 7B, S8**), T_reg_ induction (**Fig. 7C, S9**), T_reg_:CD8 ratio(**Fig. 7D**), PD-1 induction in CD8^+^ T cells(**Fig. 7E, G**), and MDSC cells(**Fig.7F**). However, coinciding with IFN-γ restoration in CD8 T cells(**Fig. 7E, G**), mice showed significantly slower progression of dysplasia and lower numbers and sizes of OSCC tumors(**Fig. 7H, I**). Based on the phenotype of Dectin-1 KO mice, dectin-1 signaling appears to play a clear role in OSCC development. Taken together, these results show dectin-1 signaling is critical for establishing an immunosuppressive milieu and the development of oral tumorigenesis.

**Fig. 7.**
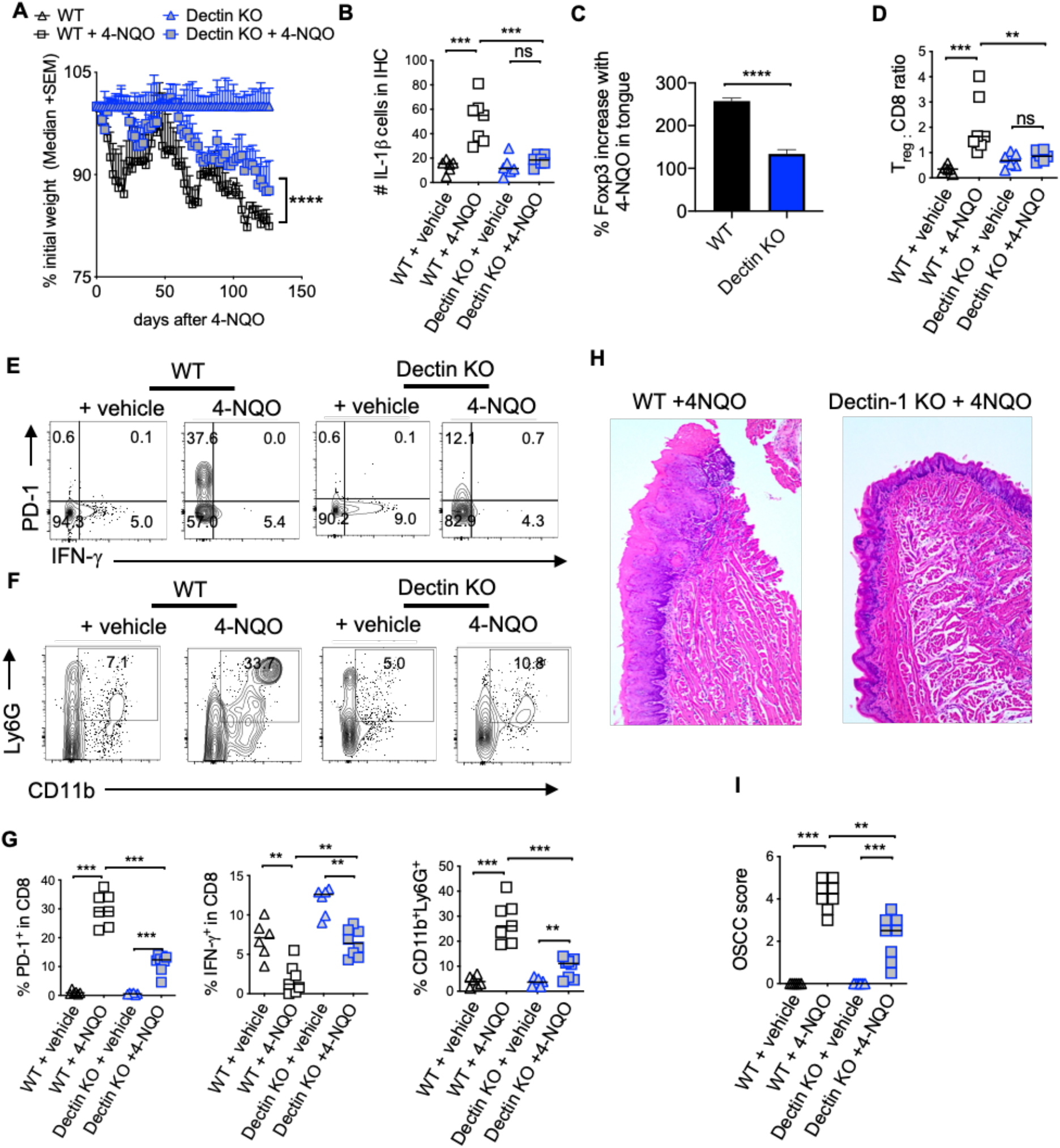
Dectin-1 deficiency reduces immunosuppressive cells and OSCC tumor burden. Statistics showing the body weight loss, comparing the indicated groups (**A**). Tongue tissues were processed for immunohistochemistry (IHC) and flow cytometry at 23 weeks after 4-NQO administration. Statistics showing the number of IL-1β+ cells in the entire field of 200X IHC image (**B**). Flow cytometry-based statistics showing % increase in T_regs_ in 4-NQO treated mice compared to Veh. controls (**C**), T_reg_:CD8 ratio (**D**), the % of PD-1 and IFN-γ expressing CD8 cells (**E**), the % of CD11b and Ly6G expressing CD45^+^ cells (**F**), and the statistical analysis for all of the above (**G**). H&E immunohistochemistry of tongue (**H**), and blinded OSCC scoring (**I**). (Mann-Whitney test except for (**B**), where t-test with Welch’s correction was performed, * P<0.05).

## Discussion

Aging skews host CD4^+^ T cells and T_regs_ towards an inflammatory T_reg_ type, which fails to control immunopathology during an infection. Instead, there is an increased accumulation of Foxp3^+^ cells, a proportion of which is associated with increased CD4^+^ T cell hyperactivation and altered levels of IL-6 and IL-1β in mice and humans in oral mucosa *in vivo* [26]. To our knowledge, no study has examined T_reg_ interactions in TIME with relevance to aging and OSCC carcinogenesis. While there are contentious data on age-related changes in T_regs_ in blood [59; 60], there is no report to date on intra-tumoral T_regs_ and their interactions with other immune suppressive cells in the context of aging. In this study, we found that mouse and human OSCC tissues showed not only an increased accumulation of T_regs_ but also of MDSC. Moreover, aged mice showed increased and early filtration of these cells and were also more susceptible to early dysplasia and tumor development compared to young mice. The 4-NQO mouse model mimicked the multistage carcinogenesis of human OSCC so that we could investigate the immunological events in tumors and tumor-draining lymph nodes in the early stages [61; 62]. In this model, we found that infiltration of immunosuppressive cells was one of the early immunological events of OSCC initiation. We also validated these findings in developed human OSCC tumors, comparing them to adjacent normal control tissues. These data provided a compelling link among aging, immunosuppressive milieu, and oral cancer. While many previous studies have shown that high levels of intra-tumoral T_regs_ associate with poor prognosis, a few studies show that intra-tumoral T_regs_ in some cancers correlate with improved disease outcomes[63–66]. However, multivariate analysis shows that T_reg_ levels are not independently prognostic of OSCC but cytolytic T cells that co-infiltrate along with T_regs_ appear to drive the favorable prognosis [15]. Therefore we examined FOXP3^+^ cells in relation to CD8^+^T cell frequency and MDSC. Increased intra-tumoral T_reg_: CD8 ratio rather than increases in T_reg_ frequencies is more informative about pro-tumorigenic functions of T_regs_ during OSCC. Indeed, examining T_reg_: CD8 ratio has allowed investigators to obtain consistent data showing clearly that a higher ratio is associated with poor prognosis and survival in human studies [67; 68]. While we also characterized the T_regs_ by examining their inflammatory and tissue markers during early carcinogenesis in aging mice, more studies are required to determine the mechanisms underlying their altered phenotype and their contribution to OSCC development.

IL-1β is a cancer-associated pro-inflammatory cytokine that is abundant in breast, colon, lung, esophageal, as well as OSCC cancers[69]. It is linked to poor prognosis for patients with esophageal cancer[70]. Targeting IL-1β has been shown to hinder oral carcinogenesis in the 4-NQO model, and is considered in the clinical settings[29; 71]. IL-1β is also involved in inducing immunomodulatory Foxp3^+^ROR-γt^+^ cells in microbiome dependent manner[26; 29; 36; 37; 51; 72–79]. Foxp3^+^ROR-γt^+^ cells have been shown to be present in tumors and contribute to tumor immune evasion and autoimmunity control[28; 80]. But the events preceding excessive IL-1β expression and at the IL-1β/T_reg_ interface are largely unknown. Our data showed that there was excessive IL-1β production in tongues of aged 4-NQO treated mice compared to young mice, demonstrating that IL-1β is clearly associated with the establishment of a worse immunosuppressive milieu. These results are distinct from an acute infection milieu and steady state-conditions where aging oral mucosa shows IL-1β suppression [26]. Although it is conceivable that early carcinogenesis environment triggers elevated levels of IL-1β in aged mice, the sequence of events that promote elevated levels of IL-1β, T_reg_ accrual, and MDSC increase during aging need to be addressed. Furthermore, we found increased dectin-1 expression in immune infiltrates in aging mice. Also, signaling through dectin-1 promoted T_reg_ and MDSC accrual. While it is clear that immune cells show activation of Syk signaling, the precise subset of cells that responds through dectin-1 signaling hyperactivation in the context of aging remains to be addressed. While we hypothesize that changes in fungal stimulation emanating from the commensal fungal microbome could lead to dectin-1 activation, this possibility remains to be further confirmed in the future. The microbiome is a significant contributor to OSCC in the 4-NQO mediated carcinogenesis model [81; 82]. Aging is also associated with changes in the resident microbiome, reduced microbiota diversity, and an increase in certain types of bacteria and fungi, resulting in local immunological changes, chronic inflammation, and other geriatric sequelae [83–85]. *Candida* carriage and related co-morbidities are significantly more frequent in elderly individuals[86]. The role of microbiome dysbiosis in enhancing dectin-1 signaling, immunosuppressive settings, and OSCC development, and how these are dysregulated during aging are critical questions to be addressed and will be studied in future investigations. Taken together, our study sheds new light on the immune-oncological events in TIME providing some key insights into mechanisms of immune evasion. By linking MDSC and T_regs_ with a higher predisposition of aged oral mucosa to tumor development, our study also provides important insights into mechanisms of higher susceptibility of the aging population to oral cancer. By revealing the pro-tumorigenic role of dectin-1 signaling and T_regs_, our study may also provide a way to control intra-tumoral T_regs_ without affecting the peripheral T_reg_ cell repertoire.

## Author contributions

PP supervised the project, designed the study, performed experiments, analyzed data, and wrote the manuscript. QP and AT provided tumor resections from human participants under an approved protocol and contributed to discussions. NB and SJ performed the experiments and analyzed ELISA and qPCR data. PM weighed the mice and performed validation qPCRs. CQ and SJ performed microscopy of the histology slides, scored the histology data in a masked fashion, performed animal breeding, and processed tissues for flow cytometry. MG and AW read the manuscript and contributed to discussions.

## Acknowledgments

PP was supported by the CWRU School of Dental Medicine departmental startup funding and NIH Head and Neck Cancer SPORE pilot grant (P30 CA043703).

## Materials and methods

### Mice and human samples

Mouse experiments were performed following all guidelines and regulations under an approval from the CWRU Institutional Animal Care and Use Committee. Young (3-4 months of age), aged (20-24 months of age) C57BL/6 mice, TLR-2^−/−^ and Dectin-1^−/−^ mice were purchased from Jackson laboratories. Human oral tissue samples were obtained by excision (tumor resection), cytobrushing (mostly epithelial cells), or by biopsy (control tissue 2-3 cm from the tumor margin) under an approved protocol approved by the University Hospitals Cleveland Medical Center Institutional Review Board.

### 4-NQO administration in mice

4-NQO was dissolved in 1:1 solution of propylene glycol and DMSO and administrated in drinking water (50-100 μg/ml) for 16 weeks, followed by the regular drinking water for another 6 weeks. Mice were sacrificed at the end of 24-25 week 4-NQO regimen. Young mice developed OSCC tumors after 24-25 weeks and aged mice around 21-22 weeks after the start of 4-NQO administration (**Fig. 1A**). Control mice received the propylene glycol/DMSO vehicle. Mouse body weight was measured every other day until sacrifice. 5-, 12 and 15- week time-points were considered early timepoints for the evaluation of immune cell changes in the tongue. Scores were assigned using tongue immuno-histochemistry using established OSCC criteria[61]. Epithelial hyperplasia and dysplasia scores were designated at 12 weeks after the 4-NQO regimen as follows: 0 = No immune infiltrates and the presence of cohesive epithelium; 1 =Sparse immune infiltrates with mild epithelial hyperplasia; 2= Sparse immune infiltrates and moderate epithelial hyperplasia and dysplasia; 3 = Severe epithelial hyperplasia and dysplasia and moderate hyperkeratosis; 4 = Severe epithelial hyperplasia and dysplasia and hyperkeratosis; 5= Presence of invasive hyperkeratosis. OSCC scores were given at 20 or 24-25 weeks after the 4-NQO regimen as follows: 0 = No immune infiltrates and the presence of cohesive epithelium;1 =Sparse immune infiltrates with mild epithelial hyperplasia; 2= Moderate epithelial hyperplasia and dysplasia; 3 = Severe epithelial hyperplasia and dysplasia and moderate hyperkeratosis; 4 = Severe epithelial hyperplasia and dysplasia and hyperkeratosis; 5= Presence of invasive hyperkeratosis, cell nests and OSCC.

### Antibodies

Purified or fluorochrome conjugated mouse and human α-CD25 (3C7 and 7D4), CD4, IFN-γ, Foxp3, CD45, CD8, CD11C, CD38, HLADR, Phospho-Syk (Ser473), IL-1β, IL-6, Arg-1, PerCP-eFluor 710 conjugated Ly-6G Monoclonal Antibody (1A8-Ly6g), APC conjugated CD11b Monoclonal Antibody (M1/70) and dectin-1 antibodies were all purchased from Life Technologies/Thermofisher and BD-Biosciences.

### Quantitative-PCR

DNA was isolated from mouse saliva swabs using the PureLink™ Microbiome DNA Purification Kit (Invitrogen). ITS and FungiQuant qPCR primers [57; 58] (Invitrogen), and SYBR Green PCR Kit (BioBasic) were used for performing qPCR, employing the real time PCR machine (Applied Biosystems). The relative amount of fungal DNA of interest was estimated from its Ct values, which were normalized to the host β-actin DNA levels.

### Histochemistry and immunohistochemistry of proteins

For immunocytochemical hematoxylin and eosin (H&E), and immuno-histochemistry staining, tongue tissues were rinsed with PBS, fixed with 10% formalin overnight, and rehydrated in 70% ethanol overnight for paraffin embedding. This was followed by paraffin-sectioning and staining by the commercial facility (Histoserv, Inc, MD). Antibodies were used at 1-5 μg/ml concentrations.

### Flow cytometry and ELISA

Cells were re-stimulated with PMA (50 ng/ml) and Ionomycin (500 ng/ml) for 4 hours in complete RPMI-1640 (Hyclone) supplemented with 10% FCS, 100 U/ml penicillin, 100 μg/ml streptomycin, 2 mM glutamine, 10 mM HEPES, 1 mM sodium pyruvate and 50 μM β-mercaptoethanol [40]. Brefeldin-A (10 μg/ml) added in last 2 hours before harvesting the cells and supernatants. For single-cell flow cytometry staining, cells washed in PBS or PBS/BSA before surface and intracellular staining. Live-Dead viability staining was used to remove dead cells in the flow cytometry analyses. Appropriate un-stain, isotype, secondary antibody, single stain and FMO controls were used. For p-Syk staining, the cells were washed, fixed and were stained with Phosflow staining kit from BD Biosciences using manufacturer’s protocol. For examining T cells, leukocyte singlets, Live-dead^neg^, CD45^+^CD3^+^CD4^+^ or CD8^+^ gates were used. For examining MDSC, leukocyte singlets, Live-dead^neg^, CD45^+^CD3^neg^ gates were used. Data was acquired using BD Fortessa cytometers and were analyzed using FlowJo 9.8 or 10.5.3 software. Mouse IL-1β and IL-6 ELISA kits from Boster Bio (Pleasanton, CA) were used to assess the protein levels in the supernatants.

### Oral *Candida* infection and Zymosan application

Mice were infected with *Candida* as previously described [36; 87]. Briefly, they were sublingually applied under anesthesia by placing a 3 mm diameter cotton ball saturated with 1 × 10^7^ *Candida albicans* (SC5314) blastospores for 90 min. Mice were re-infected as indicated. We applied Zymosan sublingually in the same manner[88].

### Statistical analysis

Results represent at least two to three independent experiments. P values were calculated by Mann-Whitney test in Prism 6.1 (GraphPad Software, Inc.) assuming random distribution. Welch’s correction and student t tests were used where indicated. One or two way ANOVA analyses were also used for grouped analyses. P < 0.05* was considered significant.

## Supplemental information

Supplementary figures are attached separately.

