## Supplementary figures for "The role of dectin-1 signaling in altering tumor immune microenvironment in the context of aging"

**Fig.S1**

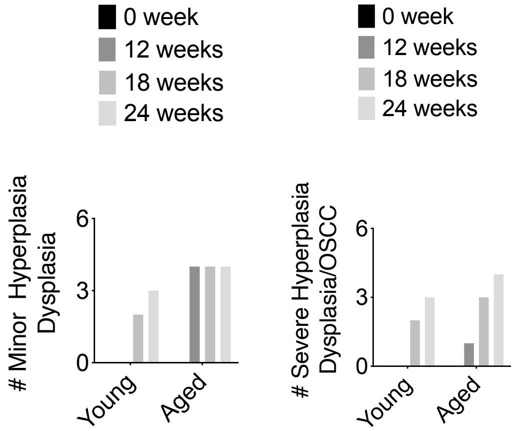


**Fig.S1. Aged mice have earlier incidence of dysplasia and more severe progression of carcinogenesis compared to young mice.** Mice were administered with 4-NQO as in Fig.1. Bar graphs showing the number of mice with hyperplasia/dysplasia in the group of 4 mice.

**Fig.S2**

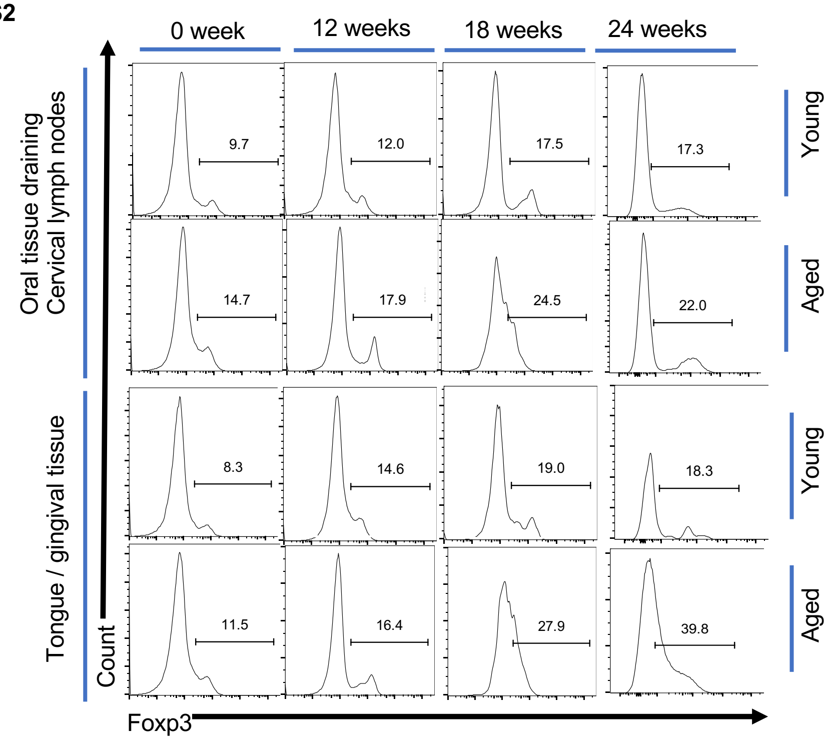


**Fig.S2. 4-NQO treatment increases the proportions of CD4$^+$T$_{regs}$ during early stages of carcinogenesis in mice. Aged mice have higher proportions of CD4$^+$T$_{regs}$ compared to young mice, before and after treatment.** 4-NQO was administered in drinking water (50 ug/ml) to mice (n=9/young or aged group) for 0,12, or 18 weeks. Samples were processed for flow cytometry. Contour plots gated on CD3$^+$CD4$^+$ lymphocyte singlets after dead cell exclusion.

**Fig.S3**

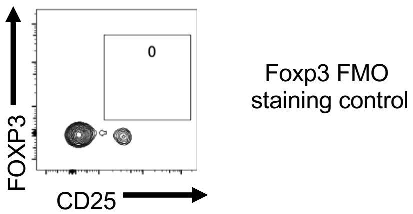


**Fig. S3. FOXP3 FMO staining.** Human oral tissue samples were obtained, processed for flow cytometry and stained with all the antibodies in the panel except FOXP3. Contour plots gated on CD3$^+$CD4$^+$ lymphocyte singlet cells after dead cell exclusion.

**Fig.S4**

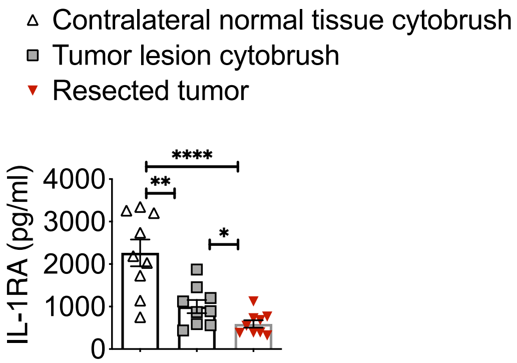


**Fig.S4. IL-1RA levels are diminished in OSCC tumors.** Human oral tissue samples were obtained either by cytobrushing or by excision under an approved IRB protocol. The single cell suspensions were restimulated with PMA/Ionomycin for 4 hours and cell supernatants were used for IL-1RA ELISA.

**Fig.S5**

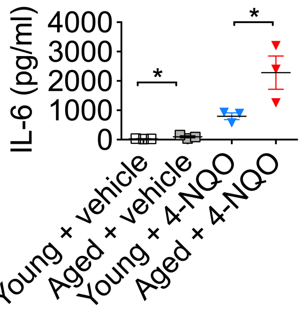


**Fig.S5. Aged mice have significantly elevated levels of IL-6 in early stages of carcinogenesis compared to young mice.** Mice were administered with 4-NQO for 12 weeks and tongue cells were re-stimulated with PMA/Ionomycin for 4 hours before supernatants from these cultures were used for IL-6 ELISA.

**Fig.S6**

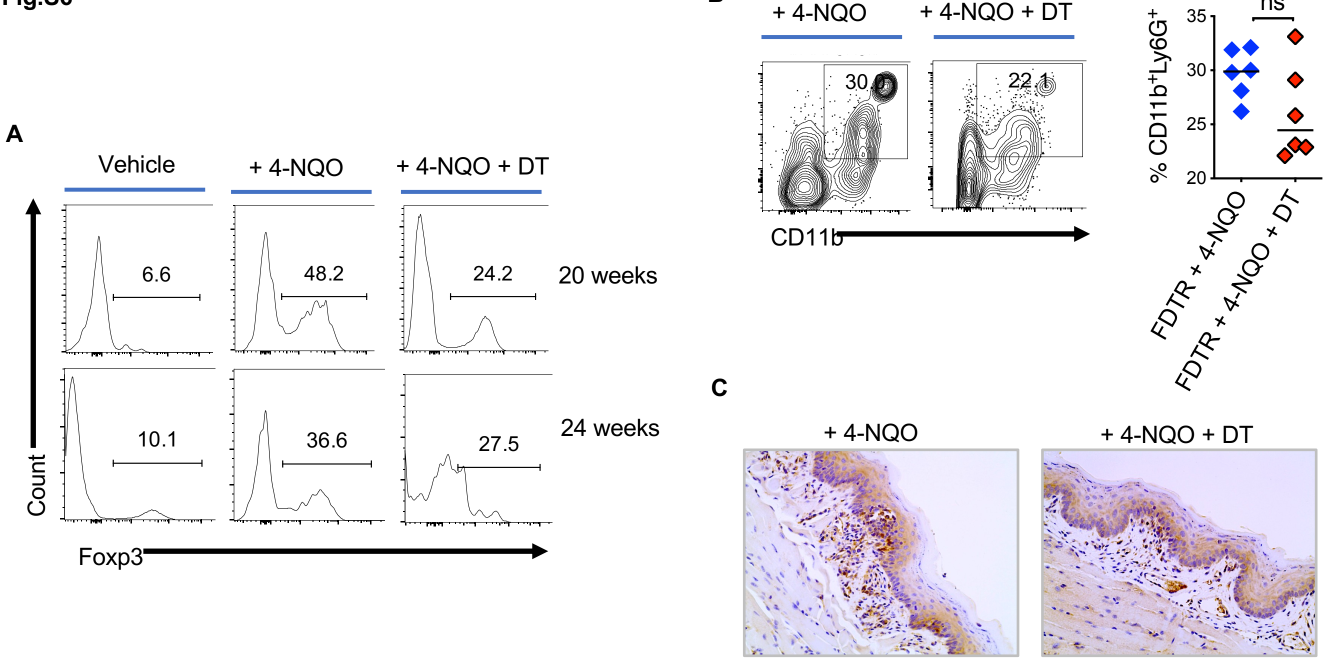


**Fig.S6. Partial $T_{reg}$ depletion does not affect IL-1β and MDSC proportions.** $T_{regs}$ were depleted in FDTR mice by injecting diphtheria toxin (DT) every 5 days between $16^{th}$ -$21^{st}$ weeks of 4-NQO treatment. Flow cytometry contour plots showing $CD4^+Foxp3^+$cells (**A**) and MDSC (**B,** left). Statistical analysis of MDSC proportions (18 weeks) (**B**, right). **C**) IL-1β immunohistochemistry staining and microscopy were performed at 18 weeks of 4-NQO administration (400X magnification).

**Fig.S7**

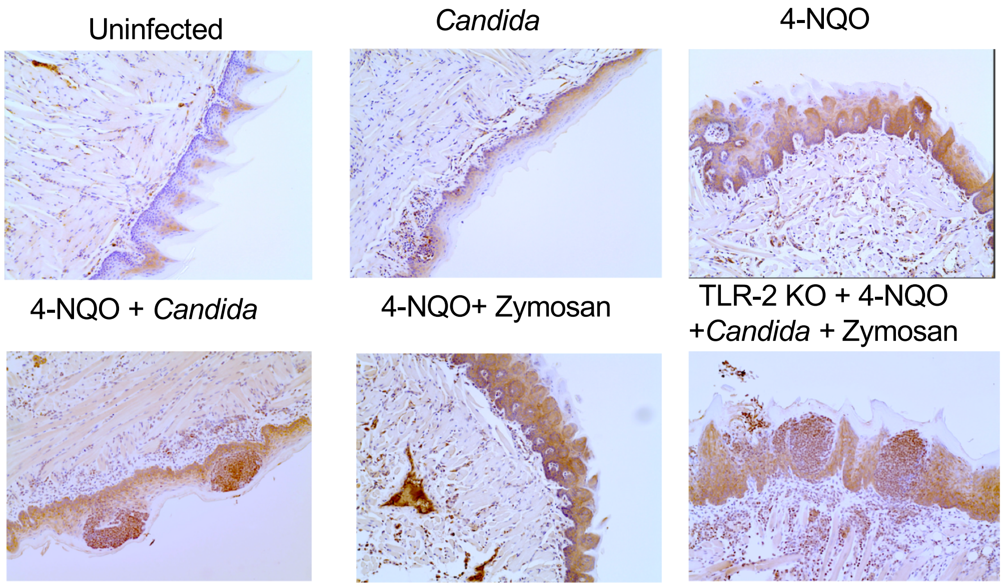


**Fig.S7. *Candida* and Zymosan increase IL-1β expression in tongue during 4-NQO induced carcinogenesis.** 4-NQO was administered in WT or TLR-2 KO C57BL/6 mice (6 mice/group). Zymosan (1mg) or *Candida* ($10^7$ blastospores) were applied sublingually under anesthesia every week between $4^{th}$ and $8^{th}$ weeks of 4-NQO administration. IL-1β immunohistochemistry staining and microscopy were performed at 12 weeks (200X magnification).

**Fig.S8**

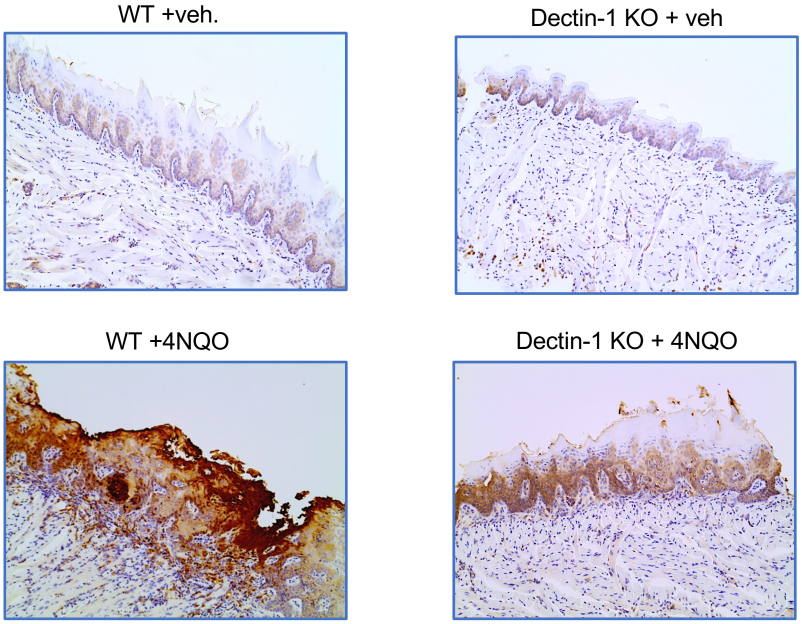


**Fig.S8. Dectin-1 deficiency reduces IL-1β expression in tongue during 4-NQO induced carcinogenesis.** Tongue tissues were processed for immunohistochemistry (IHC) and flow cytometry at 23 weeks after 4-NQO administration (200X magnification).

**Fig.S9**

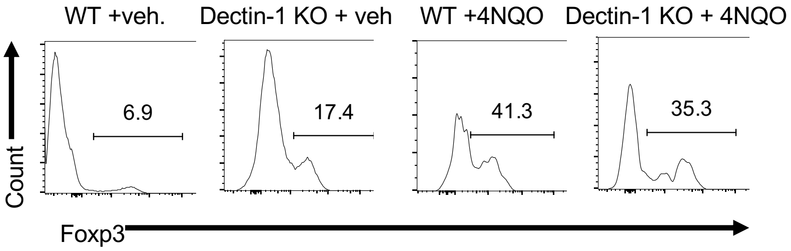


**Fig.S9. 4-NQO carcinogenesis induced $T_{reg}$ infiltration in tongue is significantly reduced with loss of Dectin-1.** Flow cytometry contour plots showing $CD4^{+}Foxp3^{+}$cells, 23 weeks after 4-NQO administration.
